# Improving the interpretability of species distribution models by using local approximations

**DOI:** 10.1101/454991

**Authors:** Boyan Angelov

## Abstract

Species Distribution Models (SDMs) are used to generate maps of realised and potential ecological niches for a given species. As any other machine learning technique they can be seen as “black boxes”, due to a lack of interpretability. Advances in other areas of applied machine learning can be applied to remedy this problem. In this study we test a new tool relying on Local Interpretable Model-agnostic Explanations (LIME) by comparing its results of other known methods and ecological interpretations from domain experts. The findings confirm that LIME provides consistent and ecologically sound explanations of climate feature importance during the training of SDMs, and that the `sdmexplain` R package can be used with confidence.

## INTRODUCTION

In recent years an increased focus has been placed on making machine learning more interpretable (Doshi-Velez and Kim, 2017; García et al., 2009). The main reason for this is that black-box models are often not trusted by domain experts (Ribeiro et al., 2016). Moreover, the visualisation of feature importances can often yield valuable insights. Such an analysis might help a scientist decide which data points are not necessary (so that data collection can be improved), or more importantly, uncover new details about what is contributing to the study phenomenon. In the very least, it can act as a sanity check that the model is learning the right things and successfully avoids bias.

Species Distribution Modeling (SDM) is the application of machine learning on estimating the species habitat based on occurrence data and associated environmental features (temperature, humidity etc.) (Elith and Leathwick, 2009). Such models can be used to guide conservation efforts, estimate the effects of climate change and answer other environmental hypotheses (Guisan et al., 2013; Austin and Van Niel, 2011). A field of such importance for ecology can benefit greatly from becoming more explainable. Domain experts and people in the field rely on the maps produced by those models, and their trust in their accuracy can be increased if they understand more clearly how models make decisions.

Some of the most important breakthroughs in explainable machine learning include the LIME Ribeiro et al. (2016) and IML (Fails and Olsen Jr, 2003) projects^1^. For this study we chose the former, due to its more accessible API (Application Programming Interface).

Those are the motivations behind the creation of the `sdmexplain` package (Angelov, 2018a). This R package has functions that enable a user to train SDMs and understand which features are most important with various visualisations and an interactive map. In order to prove that the LIME explanations used are indeed consistent with ecological observations, an analysis of two species was performed. Additionally, the LIME results were compared to a standard method for computing feature importance in order to determine in the patterns generated are statistically similar.

## METHODS

### Study species

In order to ecologically prove the explainability calculations the two species were selected based on their relatively strong dependence on specific climate conditions. The occurence (observation) data points are shown on the map in Figure 1.

**Figure 1.**
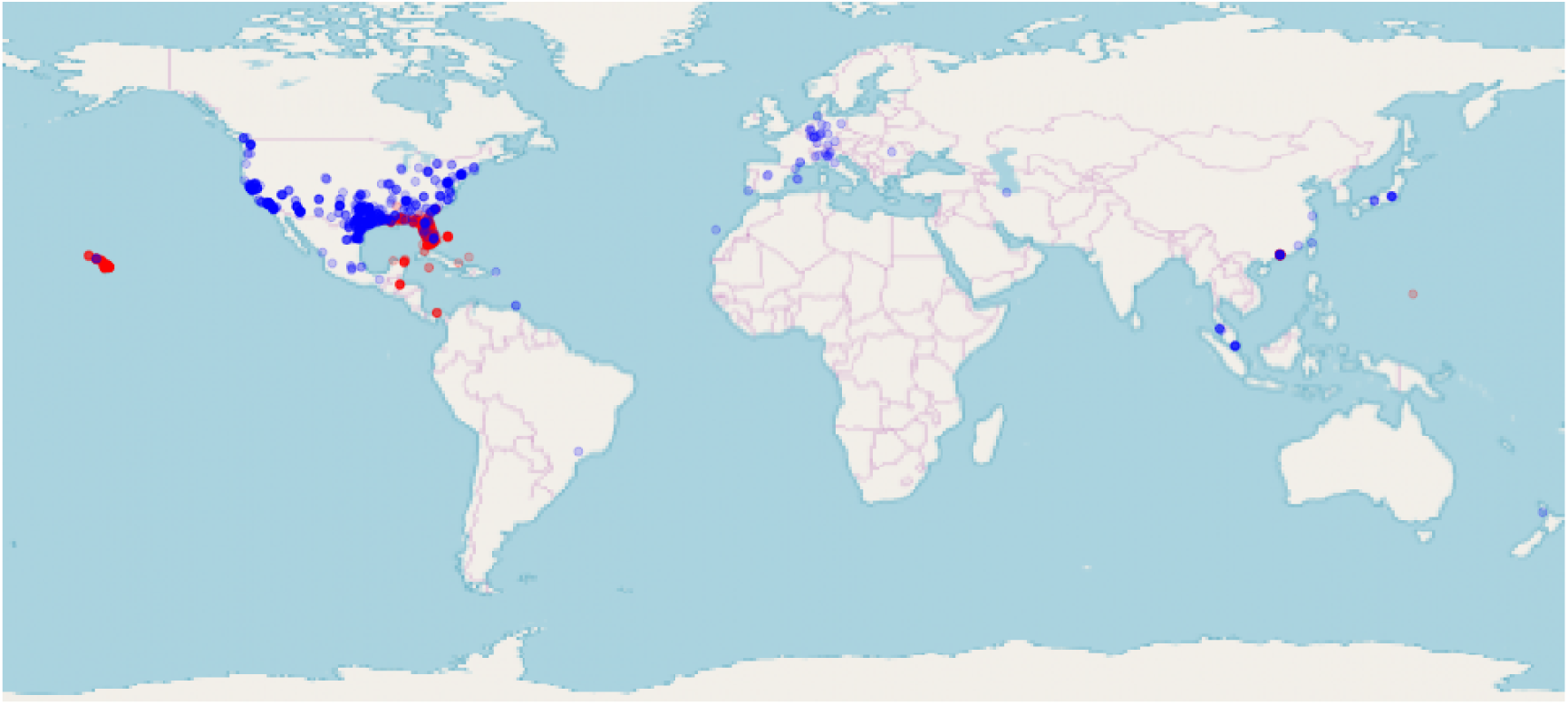
Occurence map for raw GBIF data (red: *Eleutherodactylus planirostris*, blue: *Trachemys scripta*.

### Eleutherodactylus planirostris

*Eleutherodactylus planirostris*, otherwise known as greenhouse frog, is a mostly terrestrial amphibian. It occurs natively in Bahamas, the Cayman Islands, Cuba, and invasively in Jamaica, Guam, and the southeastern United States. An important characteristic of this species is that for its eggs to hatch, 100% humidity is required (Rödder and Lötters, 2010).

### Trachemys scripta

Otherwise known as slider turtle, or common slider, *Trachemys scripta* is native to the United States, but invasive in many parts of the world, including Southern Europe, Israel, South Africa and others. Because of the negative effects of its invasive nature, this organism and its response to environmental conditions has been widely studied. The slider turtle is heavily dependent on water availability throughout the year. It’s feeding is strongly temperature-dependent. The hibernation of neonates in nests is very sensitive to low temperatures as well (Rödder et al., 2009).

### Species records

Species records have been obtained by using the `sdmbench` package (Angelov, 2018b). In the background it obtains data from GBIF (https://www.gbif.org/) and performs additional domain-specific preprocessing steps (such as removing occurences with impossible, incomplete or unlikely coordinates (based on the `scrubr` package, Chamberlain (2016)).

### Climate data

Environmental data has been obtained from WorldClim (http://www.worldclim.org/bioclim) by using the `sdmbench` package. In order to create variables that are biologically relevant, monthly temperature and rainfall data are processed. Those derived variables are more representative of seasonal and limiting climate characteristics. This preprocessing enables successful downstream species distribution modeling.

### Species distribution models

Random Forests was chosen as the algorithm to create the main species distribution model for the LIME explanations. It is a popular modeling technique that is known to perform well out-of-the box, with a good balance between efficiency, speed and high tolerance for missing data Breiman (2001). It is also one of the MLR “learners” that allow for the calculation of feature importance. The R package MLR (Bischl et al., 2016) is used because of its ease of use and accessible API. In order to compare the results from different methods, several other algorithms which support feature importance computation were also run: RF SRC (Random Forests for Survival), GBM (Gradient Boosting) and rpart (Recursive Partitioning And Regression Trees). Additionally, the Maximum Entropy (MaxEnt, Phillips et al. (2006)) algorithm was also used in the comparison. It is one of the most popular and widely-used SDM techniques, and the contribution of environmental features to the model performance can be extracted from the model.

Feature importances were compared between the MLR models, MaxEnt and LIME. For LIME the individual predictions were gathered and the average contribution values per feature computed. For the MLR models the importances are represented by the mean decrease of the Gini impurity index, shown in Equation 1, where *p*(*i*| *j*) is the proportion of samples of class *c* for a particular node *t* of the decision tree.

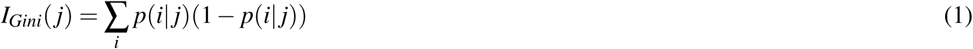

For LIME, a single explanation is computed in Equation 2, where *G* ∈ *G* is the model, *λ* (*f, g, pi*_*x*_) is the “unfaithfulness” in how the the model approximates the locality *pi*_*x*_ and Ω(*g*) is the complexity measure.

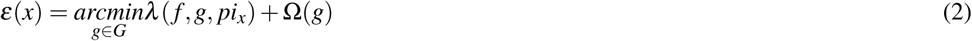

For MaxEnt, the feature importance is represented internally by the software as percent contribution. This is the result of algorithm modifying a single MaxEnt feature coefficient and assigning the gain to the environmental variable the MaxEnt feature depends on (Phillips et al., 2006).

## RESULTS

### Model Training

The data consists of the aforementioned WorldClim environmental variables and a label (or target variable). The latter contains the observations (encoded as a “positive” label). A common issue in species distribution modeling is the lack of true absences recorded (“negative” label). In those cases, in order to build a classifier, we need to artificially create those by sampling the background data, thus generating pseudo-absences (Barbet-Massin et al., 2012).

Before model training the data was randomly split into train and test sets (70% and 30% of the whole dataset respectively). This is a common method to make sure that we are not training and testing a model on the same data, which can result in overfitting (the model learns the data and its noise too well, and is prone to generealize poorly). For both species the models achieved good accuracy (% of correctly classified occurences) and Area under the curve (AUC ^2^) on the test set.

### Feature Importance Analysis

The example visualisations of LIME importances (Figure 2) show that there are two types of contributions a feature can make for a prediction: positive and negative, also with different magnitudes. Additionally, we are provided with feature rules (i.e. *bio*5 ≥ 42). Those add a new layer of interpretability to the model. A first look at those visualisations show that often a few key features are often contributing the most to the probability of occurrence at a given location. Those are quantified on Figure 3, where the mean feature importances per model are shown.

**Figure 2.**
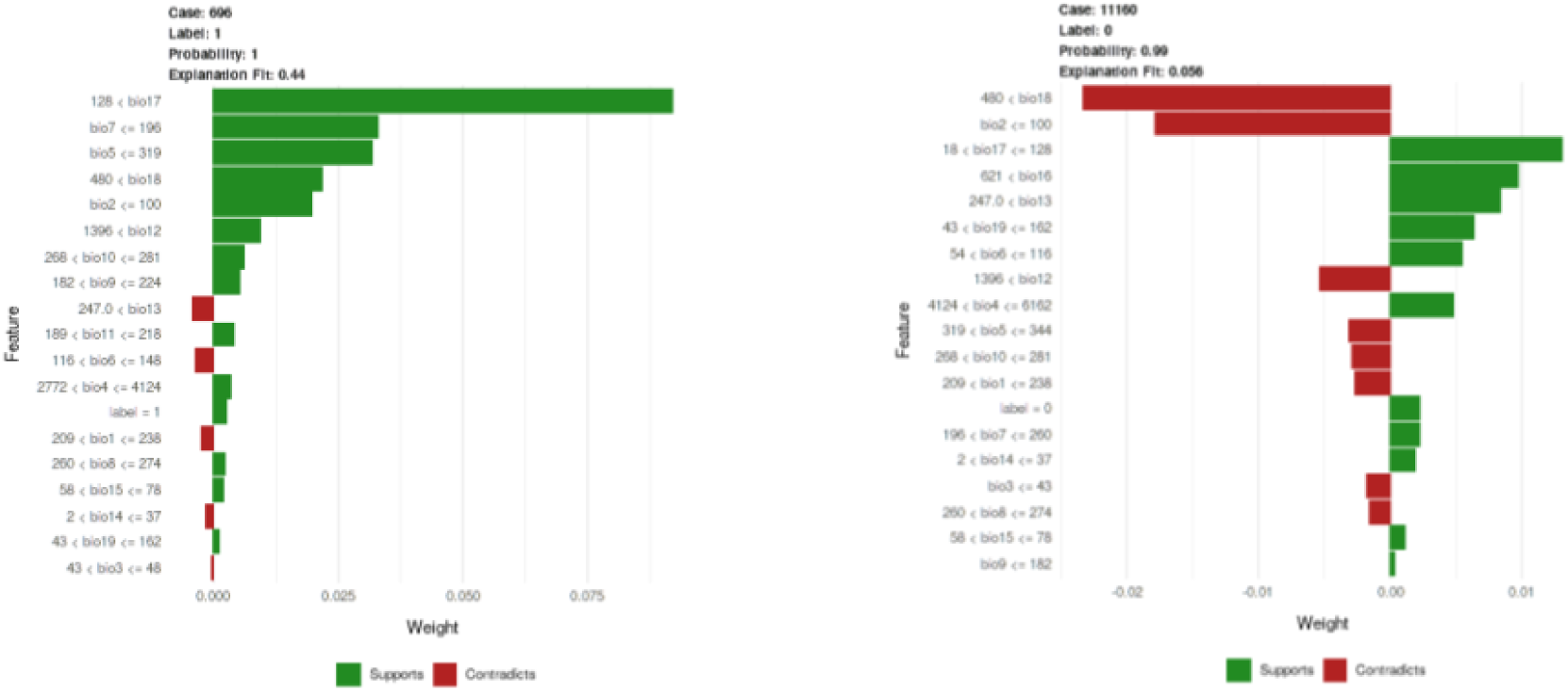
Example LIME plots.

**Figure 3.**
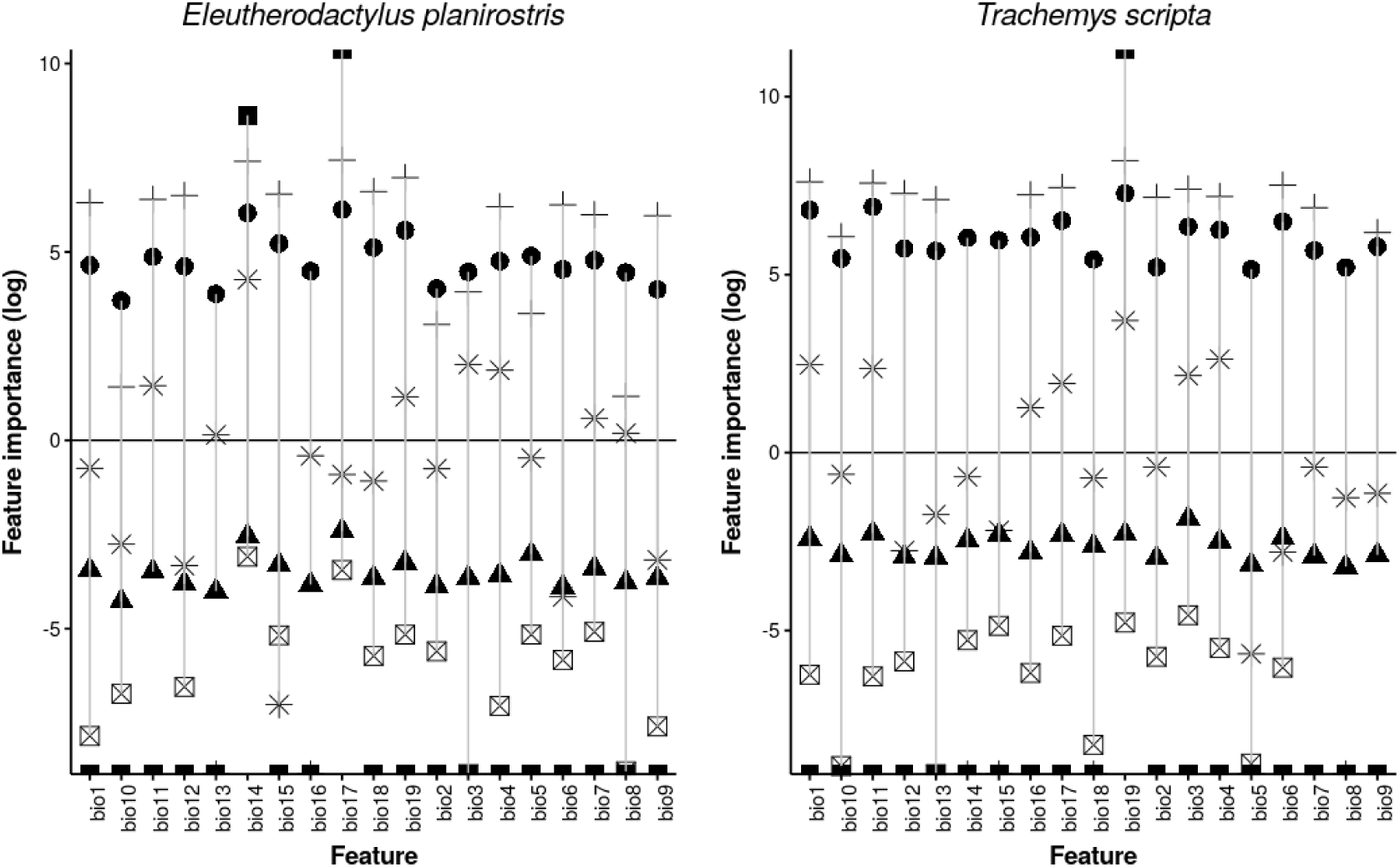
Feature importances calculated by different models (represented by shapes)

In order to test that the feature importances between the different methods are in agreement, ANOVA was performed on a multiple linear regression of species feature importance versus model type. The alternative hypothesis (that there is a significant difference between model types) was rejected with *p* > .05 for both species.

For *Eleutherodactylus planirostris* the most important features were `bio4` and `bio17`, while for *Trachemys scripta* those were `bio19`, `bio17`, and `bio3`. Those results are summarized in Table 1.

**Table 1.** Most important features for both species

| Species | Features | Description |
| --- | --- | --- |
| *Eleutherodactylus planirostris* | bio4, bio17 | Temperature Seasonality ($stdev \times 100$), Precipitation of Driest Quarter |
| *Trachemys scripta* | bio3, bio17, bio19 | Isothermality ($\frac{bio2}{bio7} \times 100$), Precipitation of Driest Quarter, Precipitation of Coldest Quarter |

## DISCUSSION

We can think of the model interpretability in three stages (Figure 4). In the first stage we are dealing with the raw data. Depending on how descriptive and intuitive the features are (i.e. the bioclimatic variables from WorldClim might be derived, but still can make sense to a domain expert), a user can already see some patterns in the data. Thus we can label this phase as a having a “medium” level of interpretability. The next step in a normal pipeline consist of the training of a SDM. This is the least understandable part of the pipeline, and most of the traditional model output is focused on various performance metrics. Some specific models, such as the ones in this study, allow for feature importance calculation (those are the various “tree-based” algorithms), but even this might not be enough, since all you get are averages across the whole dataset. Thus we can label this second phase as being the “black-box” and having a “low” interpretability. The final phase is the one that can shed a light on the black box model is the application of LIME (or similar methods) on the model, as used in the current study. Here we can achieve a very fine understanding of feature importance per observation. This can further allow for better understanding of the model mechanism, a simple sanity check that the model is learning the right things, and maybe further ecological hypothesis generation.

**Figure 4.**
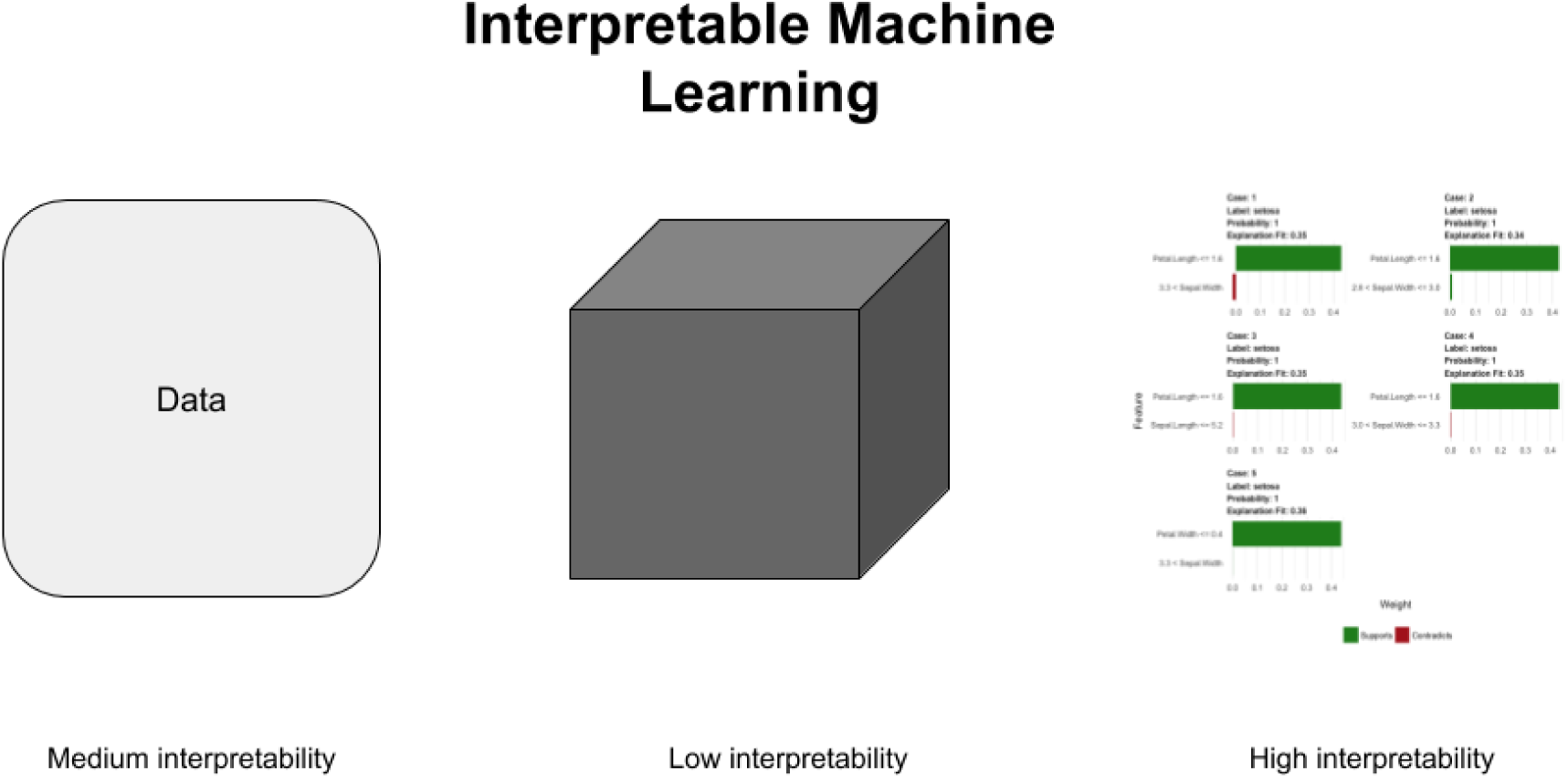
Levels of interpretability in machine learning

In statistical terms, the machine learning algorithms used are operating by using different mechanisms (despite all of them being tree based). Moreover the exact measures of “importance” also differ between the MLR methods, LIME and MaxEnt. Nevertheless, there is no statistically significant difference in results obtained across them, increasing our confidence that LIME is statistically similar.

In the case of *Eleutherodactylus planirostris*, humidity was determined to be the most important factor in limiting the species distribution. Since the hatching of the eggs requires ∼ 100% humidity this is to be expected. For *Trachemys scripta* temperature and water availability can have dramatic consequences on the behavior and reproduction of the species, and this is also confirmed by the study.

### Conclusions

This study provides scientific support for the usage of the `sdmexplain` package by confirming that LIME explanations are statistically sound and ecologically meaningful. The most important functionality of `sdmexplain` is the generation of interactive and explainable SDM maps, as shown on Figure 5. Such maps can be very useful for understanding SDMs, and providing guidance for experts in the field. Model explainability is a topic of hot research and there is a variety of new methods appearing in the field. Those can and should be evaluated in the context of species distribution modeling, especially on species of critical interest and importance.

**Figure 5.**
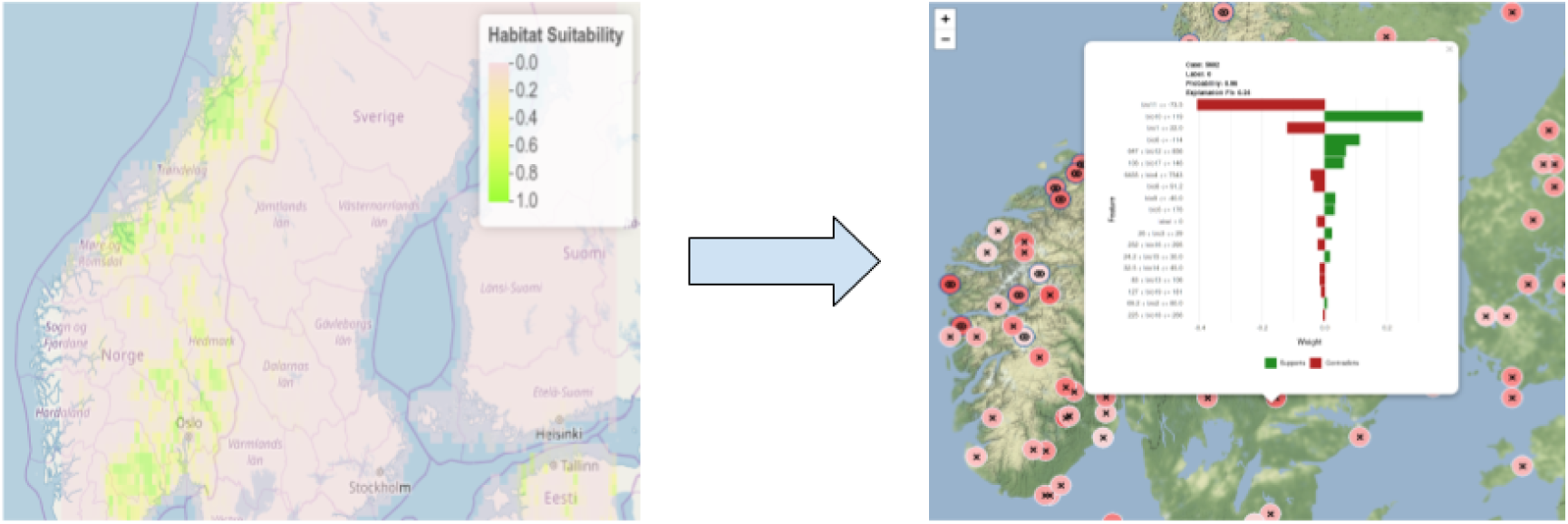
Interactive map with explainable occurences

## ACKNOWLEDGEMENTS

This research would not be possible without the original LIME package (Ribeiro et al., 2016), and its R port (Pedersen and Benesty, 2018).

## Footnotes

1 From this point onwards “interpretable” and “explainable” will be used interchangeably.

2 A standard method of estimating the performance of a binary classifier.

